# Modulation of host signalling pathways reveal a major role for Wnt signalling in the maturation of *Plasmodium falciparum* liver schizonts

**DOI:** 10.1101/2024.05.07.592925

**Authors:** Abhishek Kanyal, Geert-Jan van Gemert, Haoyu Wu, Alex van der Starre, Johannes HW de Wilt, Teun Bousema, Robert W. Sauerwein, Richard Bartfai, Annie SP Yang

## Abstract

After infection of the human host, the initial stage of the *Plasmodium falciparum* (Pf) lifecycle takes place in the liver. However, understanding of the host-parasite interaction has been limited by the rapid loss of functionality in cultured primary human hepatocytes (PHHs). Here, we link loss of hepatic functionality to drastic loss in Pf permissiveness, which we effectively prevent by using a novel medium containing serum-replacement and signal transduction inhibitors. Integrating transcriptomic analysis and phenotypic assessment of infection outcome, we identified several host signalling pathways that influence Pf liver stage development. Inhibition of the Wnt pathway in particular plays a major role in determining the size and maturity of Pf-liver schizonts, via retaining metabolic activity and epithelial nature of hepatocytes. Host signalling pathways determining Pf liver stage permissiveness provide insight into the complex host-parasite interaction and may accelerate development of novel therapeutic strategies for Pf-liver stages. (145)

## Introduction

The mosquito-borne disease, malaria, remains a devastating global burden with approximately 250 million cases and 620,000 deaths annually [1]. Most of the deaths are due to the parasite *Plasmodium falciparum* (Pf), which begins its developmental life-cycle inside liver cells (hepatocytes) after initial deposition into the skin by an infected mosquito. Over a period of approximately one week, a small number of these parasites (sporozoites) will replicate and differentiate into intracellular schizont, each containing approximately 90000 blood-infective daughter merozoites [2]. The release of these merozoites into the circulation and their subsequent cyclical asexual replication in red blood cells (RBCs) is responsible for clinical symptoms associated with the disease.

Despite its fundamental importance in enabling effective infection, currently fundamental understanding of the complex parasite-host interaction within the infected hepatocytes remains elusive. Studies with the rodent *Plasmodium berghei* (Pb) model have been informative as a proxy for Pf but findings are often not translatable to Pf-host interactions as in particular the rodent parasite has only a 2-day liver-stage period compared to 7-days for Pf [3]. Specific host membrane receptors have been identified to be involved in Pb sporozoite invasion of hepatocytes such as CD81 and Scavenger Receptor B1 (SR-B1) [4–6]; similarly, host factors involved in nutrient acquisition [7–10] and/or prevention of host cell death pathways [11–14].

Research in Pf primarily involves *in vivo* models such as mice with humanised liver [2, 15, 16] or *in vitro* models where hepatocyte cell-lines such as HC-04 [17, 18] and primary human hepatocytes (PHHs) are used. The usage of cell-lines has been limited to the identification of host receptors involved in parasite invasion such as EphA2 [19] and glypican 3 [18]. Contrastingly, PHHs are considered the gold standard in drug assays [3, 20] and used to identify host factors needed for the development of Pf liver schizont such as SR-B1 [10, 21, 22] and glutamine synthetase [23]. A well-known phenomenon of *in vitro* cultured PHHs, however, is the quick loss of hepatic features [24, 25], yet its impact has not been explored in Pf liver stages and may hinder the discovery of host factors that are important for full maturation of liver schizonts. Establishment of an *in vitro* hepatocyte model that preserve hepatocyte features to allow for late-stage Pf maturation is therefore a prerogative and the next logical frontier in research. The loss of functional hepatic features is controlled by specific host signalling pathways [24] of which, their potential impact on parasite development have not been previously examined.

Here, we set out to 1) identify the impact of loss of hepatic functions in *in vitro* hepatocytes and their permissiveness to Pf infection, 2) improve and stabilize hepatocyte culture conditions for Pf permissiveness, facilitating complete schizont development and 3) identify host pathways in hepatocytes that influence the developmental kinetics of Pf liver stages. Using RNAseq to compare the transcriptomics profile between the freshly isolated and cultured PHH, we identified upregulation of the Wnt (Wingless and Int 1), TGF-β (Transforming Growth Factor Beta), Notch and Rho signalling pathways. Treatment with specific chemical antagonists of some pathways led to improvement in Pf schizont development for some pathways. Furthermore, the Wnt pathway was shown to be most influential in determining the size of the growing Pf schizont.

## Results

### In vitro cultured PHHs lose permissibility for Pf liver stage infection and development

Traditionally fresh primary human hepatocytes (PHH) are cultured in standard medium containing human sera (WBH) [23, 26, 27] and infected with Pf at 48 hours after plating. However, cultured hepatocytes quickly lose their cellular identity, and this may impact on the host cells’ permissibility to sustain Pf infection. To investigate the effect of longer culturing, we infected PHH at different time points post-plating (p.p) with different Pf strains with different developmental kinetics (PfNF54, PfNF175 and PfNF135 [23]); schizont numbers were evaluated at five days post-infection (p.i) (Figure 1A). We observed a sharp decrease in number of schizonts in relation to the p.p. period for all Pf strains tested with the most drastic decline occurring between days 2 and 5 p.p. This reduction in the number of infected cells could not be explained by a decrease in the number of viable hepatocytes, which became prominent only after day 9 p.p (Figure 1C).

**Figure 1:**
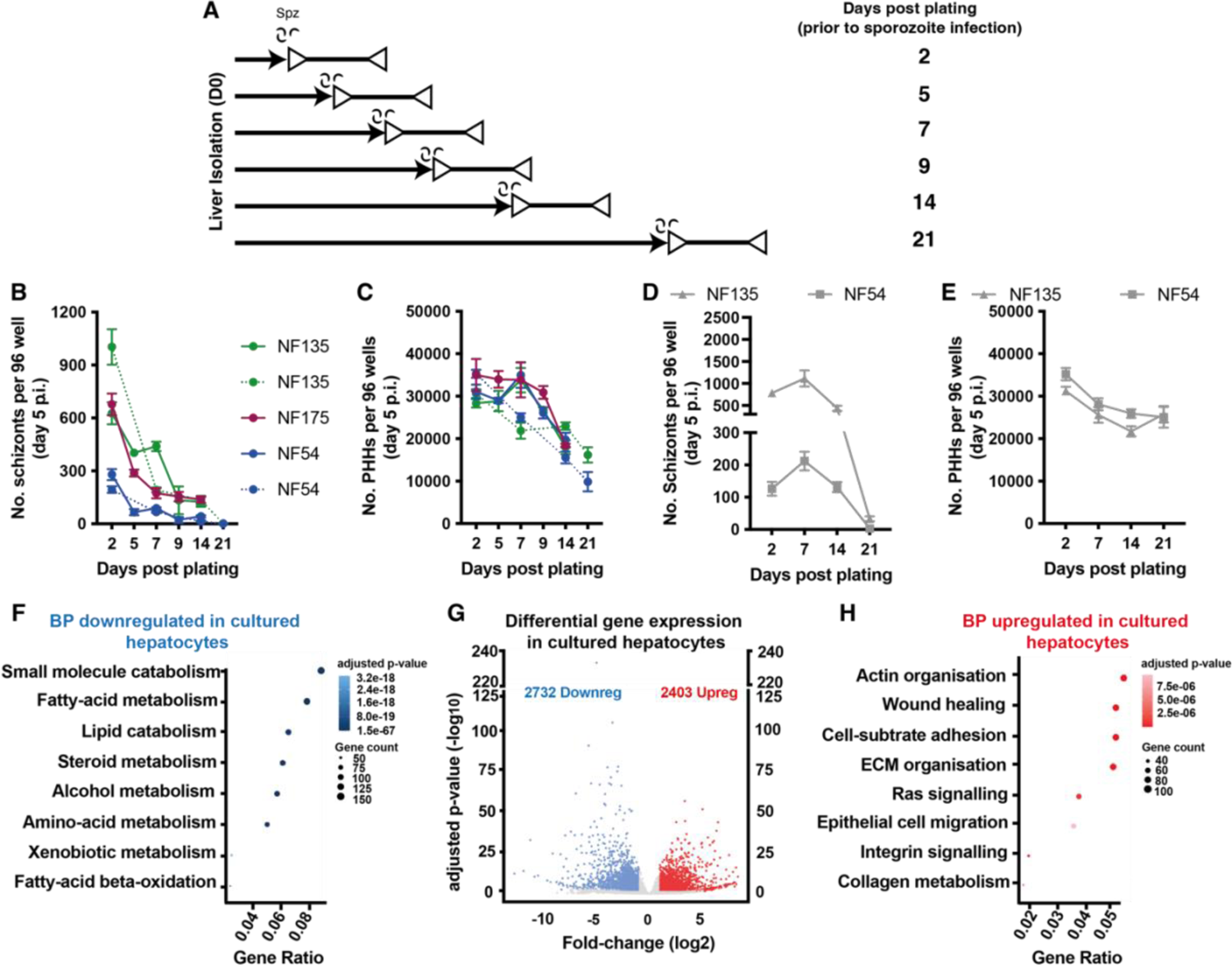
*Pf-*schizont development post-hepatocyte plating and transcriptomic profiles of primary human hepatocytes (PHH). A) The solid arrows show the number of days after liver isolation (post-plating) and the lines with the clear triangles show the period of schizont development. B) The number of schizonts in PHH on day 5 post infection (p.i.) when infected on different days p.p. for parasite strains: NF175 (magenta;), NF135 (green;) and NF54 (blue) in two independent experiments (except NF175). C) The number of PHHs present on day 5 p.i. on indicated days p.p for the specific parasite strains. For each independent experiment, the mean of three replicates is shown along with the standard deviation. The number of schizonts (D) and hepatocytes (E) on day 5 post invasion (p.i.) for NF135 and NF54 (n=1) for different infection times post plating (p.p.) cultured in B27 medium (grey). Dot-plots representing GO terms for biological processes of (F) downregulated and (H) upregulated in cultured hepatocytes. (G) Volcano plot representing the differential gene expression in cultured hepatocytes.

A previous study has shown that medium containing human sera drastically affects function of hepatocytes [21]. Therefore, we examined the impact of sera on hepatocyte permissiveness for parasite development via the removal of sera (WB versus WBH) (Figure S1). Only mild improvements in schizont numbers were observed in sera negative medium (WB) as well as presence of the maturation marker PfMSP1. We hence concluded that a novel hepatic culture medium is needed to return schizont numbers to those of day 2 p.p. PHH. A serum replacement supplement (B27) on a William’s E Glutamax base was tested given its beneficial effect for hepatic differentiation of stem cells [28] and hepatic organoids [29, 30] as well as other hepatic culture systems [24]. Addition of B27 to into the basal medium William’s E Glutamax substantially delayed the loss of hepatocyte permissiveness by at least 7 days (Figure 1D). However, this was not due to changes in hepatocyte numbers which remained comparable to the WBH medium thus reflecting a change in the phenotype of the PHHs (Figure 1E).

Irrespective of these positive effects on Pf infections, uninfected PHHs cultured for 7 days displayed a transcriptional profile that was vastly different from freshly isolated PHH showing an upregulation of 2403 gene transcripts and a down regulation of 2732 genes (Figure 1G). The most significantly up-regulated genes were those involved in actin filament organization, cell-substrate adhesions, wound healing and extracellular matrix organization (Figure 1H). All these pathways are classically associated with epithelial to mesenchymal transition (EMT) [3]. As for down-regulated genes, we found a strong enrichment for those involved in metabolism i.e. small molecule catabolism, fatty acid metabolism, organic acid catabolism (Figure 1F). Together, while serum replacement (via B27) has a substantial positive effect on PHH permissiveness to Pf infection, it cannot fully prevent the loss of hepatic functions, which may be potentially relevant for the maturation of liver schizonts.

### Specific host signal transduction pathways influence Pf liver stage development

Testing the potential relevance of specific signalling pathways in culture-induced upregulation of genes (Figure 1F) showed a strong enrichment of genes associated with Wnt (Wingless and Int-1) signalling, Ras protein signalling, transforming growth factor beta (TGF-β), intrinsic apoptotic signalling pathway and Notch signalling pathways (Figure 2A). Compared to the fresh PHH, genes involved in the Wnt, TGF-β and Notch pathways were strongly upregulated (Figure 2B-D), while BMP and Adenylyl-cyclase pathways mildly upregulated in B27 cultured PHH (Figure 2E). To examine the impact of these host pathway on Pf liver stage development, we treated PHH for 6-8 days p.p., with selective modulators of the individual pathways (IWP2, SB431542, DAPT, LDN193189 and Forskolin) (Figure 2F-J). Quantitative measurements assessed the replication (size, nuclear DNA content via DAPI; Figure 2F-G), health (human glutamine synthetase: hGS; Figure S2B-C) and maturation (PfMSP1; Figure 2I) status of the schizonts, respectively (Figure 2F-J, Figure S3). Modulation of both the BMP and the adenylyl-cyclase pathways through the usage of LDN193189 and Forskolin, respectively did not result in significant improvements in neither the number of schizonts (Figure 2F) nor phenotypic parameters. DAPT-treated hepatocytes contained parasites displayed limited, but significant improvements in the number and the size of the schizonts. Improvements were found by SB431542 treatment for all five parameters most notably in number of schizonts and expression of maturation marker, PfMSP1. Similarly, in IWP2-treated hepatocytes schizonts showed improvements especially in size, DNA content and expression of PfMSP1. Finally, we tested a previously published cocktail containing all five of these inhibitors (5C) [24] (Figure S3). Interestingly, hepatocyte monolayers cultured in the presence of 5C showed poor support of PfNF175 schizont size and numbers (Figure S4D-J). Together, these data demonstrated that inhibition of specific host pathways, and in particular Wnt, TGF-β, are important for supporting Pf permissiveness and/or schizont development.

**Figure 2:**
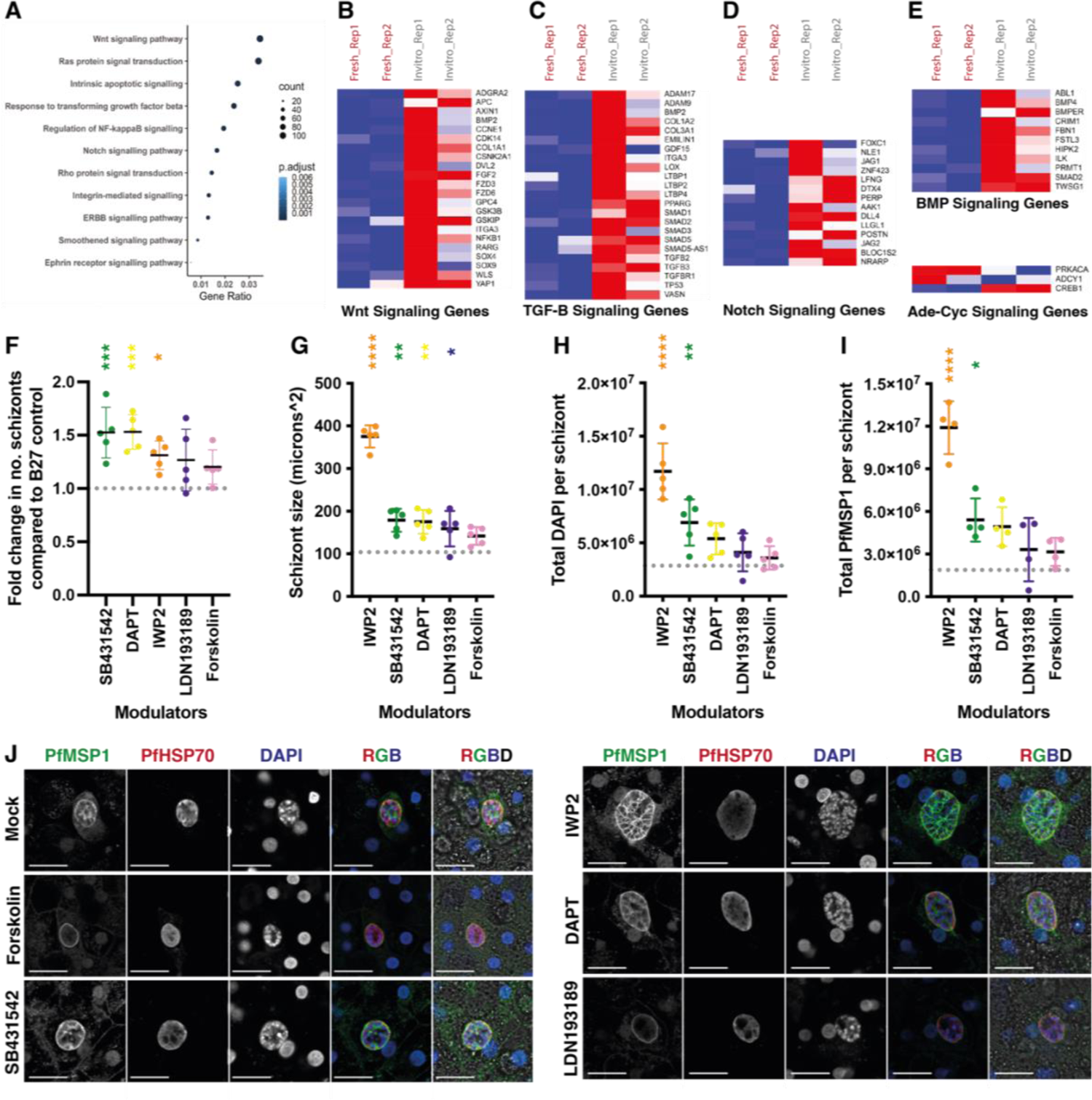
Effect of signalling pathways on Pf schizont development. A) Dot plots representing pathways that are upregulated in hepatocytes cultured in B27 for 7 days post plating. Heatmaps representing normalized expression of genes from (B)Wnt, (C)TGFβ, (D)Notch, (E)BMP and Adenylate cyclase signalling pathways deregulated in cultured vs freshly isolated hepatocytes. The data is scaled and representative of 2 biological replicates of RNA sequencing. The effect of specific host pathway inhibitors (IWP2, SB431542, DAPT, LDN193189 and Forskolin) on respectively schizont numbers (F; each dot is the average count of two technical replicates), size of schizonts (G; each dot is the median of at least 100 schizonts), total DAPI per schizont (H; each dot is the median of at least 100 schizonts), and total PfMSP1 per schizont (I; each dot is the median of at least 100 schizonts) on day 5 p.i.. The grey dotted line (per graphs F-I) shows the median measurements of schizonts grown in B27 control (from four independent experiments). The p-values from a Dunnett’s multiple comparisons test are shown. J) Representative confocal images (from four independent experiments) showing schizonts grown in B27 alone and B27 supplemented with Forskolin, SB431542, IWP2, DAPT and LDN193189 on day 5 p.i. stained with PfMSP1, PfHSP70 and DAPI. Scale bar is 25 microns.

### Inhibition of the Wnt preserves metabolic profile and epithelial nature of PHHs and improves developmental kinetics of Pf

To study the earliest impact of Wnt pathway on Pf liver stage development, PHHs were treated with either IWP2 (6 days) prior to and during Pf infection (Figure 3A-F). IWP2 treatment promoted parasite development (size, nuclear content and PfMSP1 expression) from day 3 p.i. onwards for all Pf strains. To explain the developmental kinetics elicited by inhibition of the Wnt pathway, RNA sequencing was performed on hepatocytes cultred in either B27 or B27 supplemented with IWP2 at day 5 p.i. Based on spatial proximity in the PCA space, B27 cultured cells were very distinct from fresh hepatocytes (83% of variance, Figure 4A) as also evident from Figure 1F-H. This difference decreased upon addition of IWP2. Thus, treatments with IWP2 suppressed the transcriptional changes observed during the *in vitro* culture in B27 medium (possibly associated with loss of native tissue context). Interestingly, 5C-treatment elicited additional transcriptional changes apparently not beneficial for Pf development (Figure 3G). In IWP2-treated cells, 110 genes were upregulated and 312 genes downregulated relative to B27 cultured cells (mock, Figure 3I). Gene Onthology analysis of all the genes upregulated in IWP2 treated PHHs revealed enrichment of metabolism associated terms like fatty acid, alcohol and steroid metabolism (Figure 3J). Among the biological processes suppressed upon IWP2 treatment, we found enrichment of extra-cellular matrix organization, surface-adhesion, epithelium migration and Wnt-signalling (Figure 3H). We, hence, concluded that IWP2 treatment improves the metabolism profile of cultured PHHs while simultaneously suppressing dedifferentiation processes like epithelium to mesenchymal transition, which associates with an improved developmental kinetics of liver schizonts.

**Figure 3:**
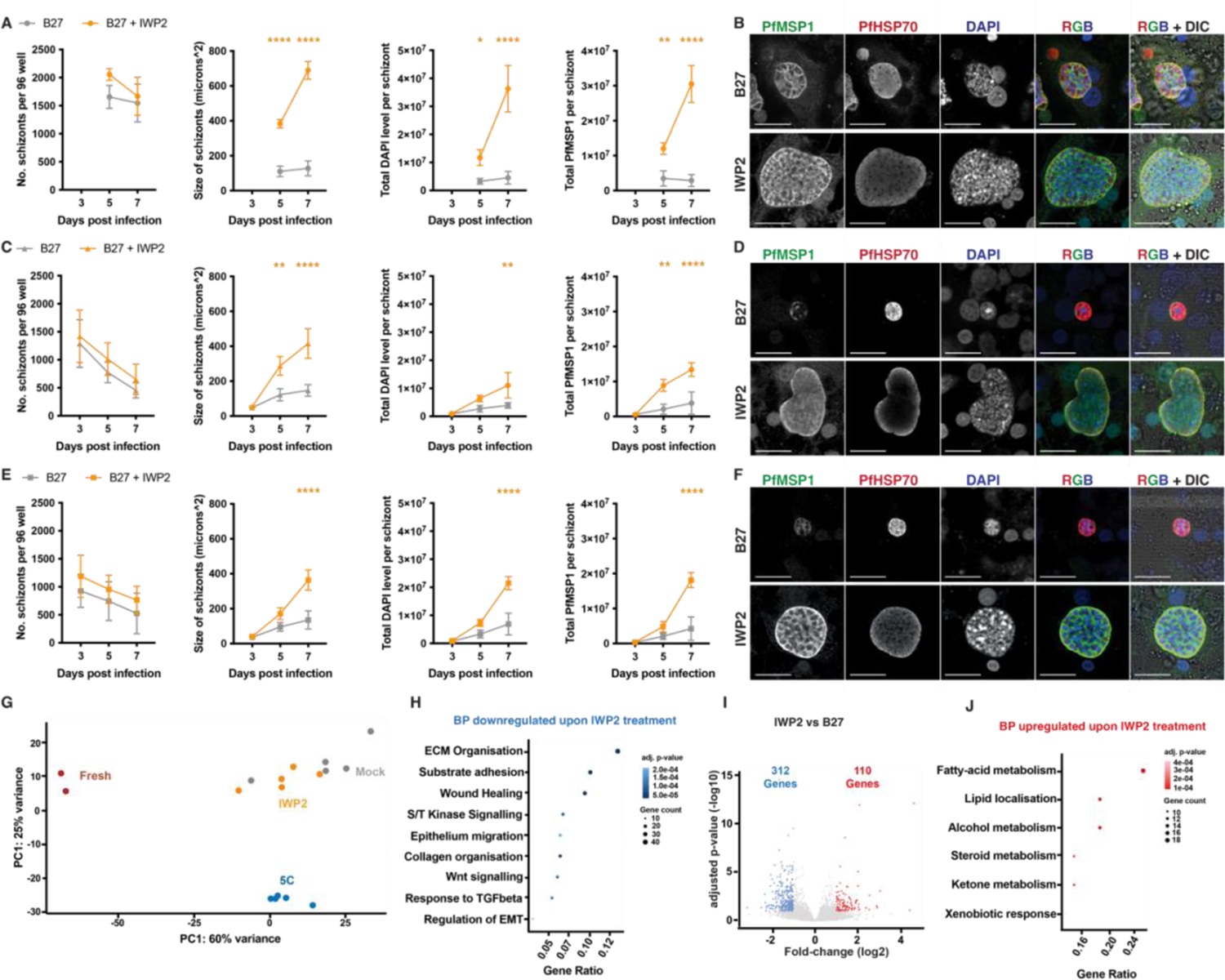
Effect of IWP2 on developmental kinetics of NF175 (A-B). NF135 (C-D) and NF54 (E-F) schizonts. For each Pf strain, the number schizonts, the size of schizonts, the total DAPI content and total PfMSP1 content are shown from left to right. The schizonts are either grown in hepatocytes cultured in either B27 (grey) or B27 supplemented with IWP2 (orange) for 6 days prior to infection and also during schizont development on days 3, 5 and 7 p.i.. Note briefly that day 3 schizonts of NF175 was not examined. For the number of schizonts, the mean of three independent experiments, each with 2-3 internal replicates is shown. For the schizont size and the total DAPI per schizont, at least 100 schizonts per each of three independent experiments (with two replicates per experiment) were measured. The mean of the median from each experiment is plotted and the graph shows the mean with standard deviation. For the total MSP1 content, at least 100 schizonts per each of three independent experiments were measured. The median is plotted and the graph shows the mean with standard deviation. A Dunnett’s multiple comparisons test is performed, and the p-values are displayed. B, D, F) Representative confocal images (from three independent experiments) showing NF175, NF135 and NF54 schizonts grown in B27 alone and B27 supplemented with IWP2 on day 7 p.i. stained with MSP1, HSP70 and DAPI respectively. Scale bar is 25 microns. (G) PCA plot representing distinct transcriptional states of freshly isolated and in vitro cultured hepatocytes under mock, IWP2 (and 5C) treatment. Dot-plots representing GO terms for biological processed (H) downregulated and (I) upregulated in IWP2 treated hepatocytes. (J) Volcano plot representing the differential gene expression in IWP2 treated hepatocytes (downregulated genes in blue and upregulated in red; fold-change (log2) >2-fold and adjusted p-value<=0.1).

**Figure 4:**
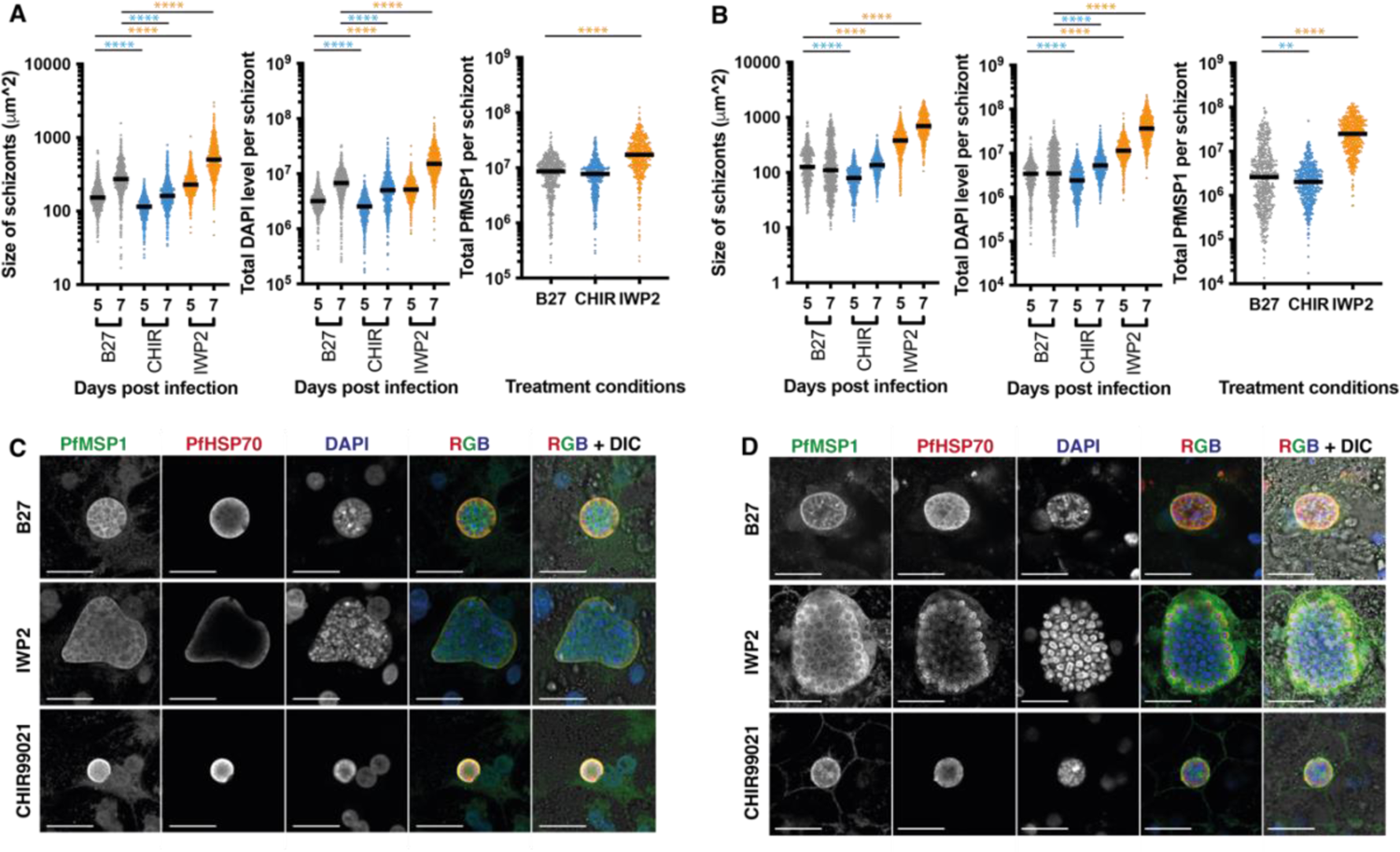
The impact of the Wnt signalling pathway on schizont size. Hepatocytes treated with B27 or B27 supplemented with IWP2/CHIR99021 and infected with either NF175 (A) or NF135 (B). Schizont size (left), and total DAPI (middle) content are measured on days 5 and 7 post infection. The total amount of PfMSP1 per schizont was measured on day 7 p.i. (right). Each dot represents a schizont with at least 100 schizonts measured for three independent experiments. The median is shown, and a Dunn’s multiple comparisons test is performed with the p values displayed. Representative confocal images (from three independent experiments) showing NF175 (C) and NF135 (D) schizonts B27 or B27 supplemented with IWP2/CHIR99021 on day 7 p.i. stained with MSP1, HSP70 and DAPI. Scale bar is 25 microns.

### Activation status of the Wnt pathway determines the size of Pf schizonts

To further examine the impact of the Wnt pathway on Pf schizont size, we treated PHHs with the Wnt activator CHIR99021 prior and during PfNF135 and PfNF175 infection (Figure 4A-D). Compared to the B27 control, the number of schizonts were mildly increased for IWP2 treated PHHs but not consistently changed for CHIR99021 (Figure S4A-B). As predicted, schizonts in CHIR99021-treated PHHs were slightly but significantly smaller than B27 cultured schizonts on day 5 p.i. for both Pf strains. There was also a significant size decrease for PfNF135 but not for PfNF175 at day 7 which may be the result of arrested development of PfNF175 under B27 conditions (Figure S3). CHIR99021 treatment also resulted in a relative decrease in PfMSP1 expression significant for PfNF175. Thus, both inhibition and activation of the Wnt pathway in hepatocytes affects predominantly schizont size and maturation. Collectively, it demonstrates that the essential role played by the Wnt signalling pathway of host hepatocytes in determining schizont size and potentially developmental kinetics.

## Discussion

In this study, we show that primary human hepatocytes in *in vitro* culture quickly lose their ability to sustain a Pf infection, irrespective of the parasite strain. This loss of permissibility coincides with up-regulation of specific signalling pathways in hepatocytes. Inhibition of individual up-regulated pathways (Wnt, TGF-β, and Notch) results in the enhanced or maintained hepatocyte permissibility and/or improvement in schizont development. In addition, we identify the Wnt-pathway as an important host factor that determines the size and maturity of the developing schizonts.

Although combined blockage of all these up-regulated pathways greatly benefits hepatitis B virus infection as shown previously [19], it appears detrimental to hepatocytes permissiveness and development of Pf as shown in this study (Figure S4). This could be due to the differing intracellular host requirements by the two distinct pathogens: for example, the intrahepatic growth of HBV is 72 hours compared to the 7 days of Pf. Furthermore, 5C treatment appear to exert transcriptional and/or cellular changes beyond the prevention of EMT (Figure 3A). In case of Pf, we show here that specific individual host pathways rather than the combination of all pathways bring strong improvement on parasite development.

We find that the inhibition of the Wnt pathway in cultured hepatocytes have a marked effect on parasite size and maturation. This increase in parasites size is concordant with the upregulation of genes involved in lipid, alcohol and steroid metabolism (Figure 4). While general metabolic health of hepatocyte is conceivably beneficial for parasite growth, fatty acid synthesis in particular is relevant to parasite development and formation of hepatic merozoites [31]. Furthermore, upon Wnt inhibition we find a downregulation of genes typical for EMT transition. In the context of *in vitro* cultures EMT is mainly due to the loss of tissue environment. EMT however can also occur *in vivo* during liver fibrosis [32] and might influence liver stage infection.

While our study highlights the relevance of the inhibition of the Wnt signaling pathway in *in vitro* cultures, this observation might also relevant during *in vivo* zonal differentiation of hepatocytes. It is well established that within the liver lobule hepatocytes display marked difference in gene expression and metabolism along the portal central axis (also refer to as zonation, [33]) and components of Wnt pathway are predominantly present in zone 3 hepatocytes and heavily involved in maintaining hepatic zonation [34]. We have previously shown that Pf parasites strongly prefer zone 3 (pericentral) hepatocytes resulting in better parasite development [23], as similarly shown for *P. berghei* [35]. While zone 3 hepatocytes possess all components of the Wnt pathway i.e. the surface receptor and intracellular signalling components, the actual external Wnt ligand/signalling proteins may be possibly secreted by hepatocytes from neighbouring zones or other non-hepatocyte cells present in the liver [36] and may therefore also play a role in determining schizont size.

The slight increase in schizont size observed with the other pathway inhibitors may possibly also relate to the Wnt status. Schizonts grown in PHHs treated with forskolin (adenylate cyclase activator) or LDN193189 (inhibitor of Bone Morphogenic Protein pathway) are the relatively smaller with lower PfMSP1 expression. In forskolin treated schizonts, activated adenylate cyclase leads to increased levels of cAMP which in turn can activate the Wnt pathway[37, 38]. However, the relationship between the BMP pathway (LDN193189) and the Wnt pathway is less obvious as the former can either inhibit or activate the Wnt pathway depending on the presence of wildtype of p53 or the loss of SMAD4 respectively [39]. This may explain the large variation observed in schizonts grown with LDN193189. Inhibition of the Notch pathway (by DAPT) maintains PHHs in their hepatocyte lineage (as opposed to a more bile ductal phenotype) as well as indirect inhibition the Wnt pathway [40]. This could hence explain the observation that, schizonts grown in DAPT-treated PHHs are larger compared to the control B27 medium. Finally, TGF-β signalling leads to the activation of the Wnt pathway [41] therefore its inhibition results in relative better schizonts as for size, nuclear content, and maturation.

Despite their differences in infectivity and phenotype, schizonts of all tested Pf strains are significantly larger and more mature in IWP2-treated PHHs than their counterparts in B27-treated PHHs. This larger size becomes significant during the process of schizont growth i.e. after day 3 p.i. and becomes prominent at day 5 p.i.; blockage of Wnt communication between infected- and uninfected cell could allow for better expansion of the growing schizont with less of host restriction. It has been estimated that an infected hepatocyte can expand up to 200 times its original volume to accommodate the growing Pf schizont [42]. This could necessitate some communication between the infected and uninfected hepatocyte and the Wnt signalling pathway may serve here as communication platform [33,35–37]. IWP2 specifically inhibits the enzyme, porcupine (Porcn), which is involved in transport of the wnt ligands [43]. One could hypothesize that neighbouring hepatocytes keep their strict size and volume via the steady basal level of wnt signalling and thereby limiting the growth of Pf schizonts [44]. A possible mechanism of action is summarised in Figure 5.

**Figure 5:**
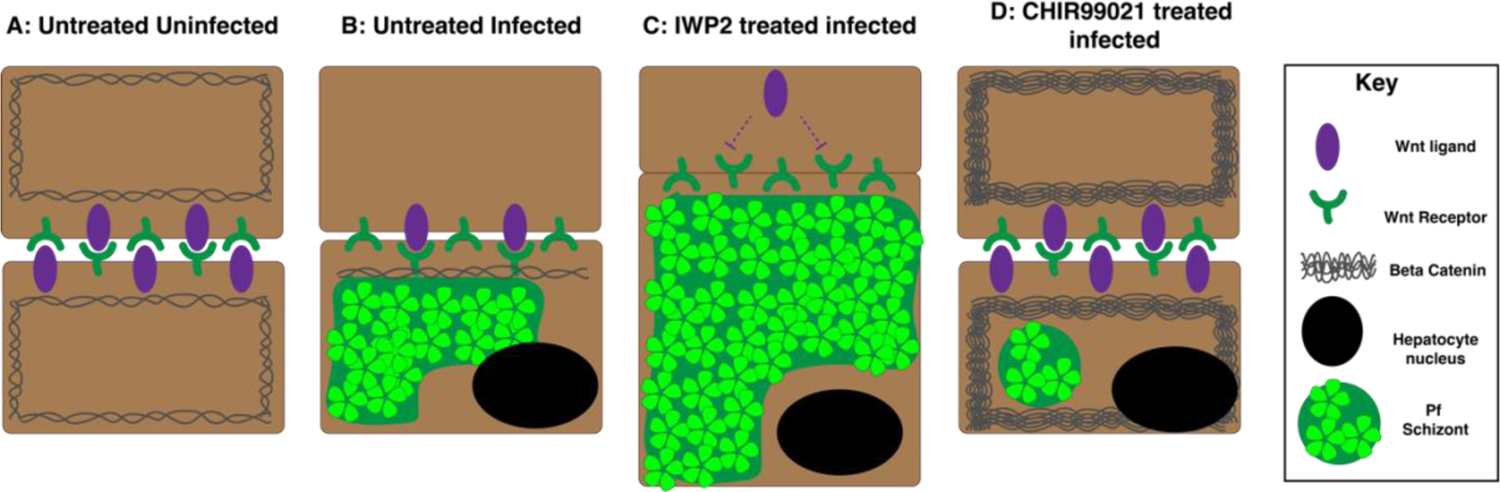
Schematic of a possible mechanism in which the Wnt pathway controls Pf schizont development. A) Under untreated and uninfected condition, there is a steady level of beta-catenin (β-catenin). In addition to its role in gene transcription, β-catenin is the intracellular component that links the cadherin adhesion molecules to actin filaments and therefore controls the “rigidity” of the cells in relation to its neighbours. B) In an infected hepatocyte (after 3 days post infection), the growing Pf schizont takes up so much of the hepatocyte volume that the trafficking of the Wnt ligands to the surface is disrupted. Wnt receptor of the neighbouring uninfected cells are not activated, and β-catenin molecules are degraded due to the phosphorylation of the enzyme, glycogen synthase kinase 3 (GSK3): this allows some flexibility in the uninfected neighbouring cells to accommodate the growth of the infected cell. However in these (uninfected cells), Wnt ligands are still present and can interact with the existing Wnt receptors on the infected hepatocytes to maintain some β-catenin in the infected cell, thus limiting the growth/size of the schizont. C) Under IWP2 treatment, Wnt ligands are not trafficked to the surface of both uninfected and infected hepatocytes due to the inhibition of the enzyme porcupine (target of IWP2). Porcupine “labels” (via palmitoylation) Wnt ligands for correct trafficking to the plasma membrane. As a result, Wnt receptors on both uninfected and infected cells are not activated and existing β-catenin are degraded leading to reduced connection between cadherin and actin filaments (i.e. cell-cell contacts) and ultimately reducing rigidity. D) Under CHIR99021 (GSK3 inhibitor) treatment, β-catenin in both uninfected and infected cells cannot be phosphorylated nor degraded. Both cell types are more rigid due to the improve connection between actin filaments and cadherin which severly limits the size of the Pf schizont.

In conclusion, we show that PHHs cultured in the standard medium (WBH) rapidly lose their permissiveness to Pf liver stage development. This loss, however, can be substantially delayed by the usage of a defined serum replacement (B27) on top of a William’s E Glutamax base. Furthermore, we identify three specific host signalling pathways (Wnt, TGF-β and Notch) to modulate Pf liver stage development. In particular, the Wnt signalling pathway is shown in a novel role to control the size of the developing Pf schizont and is therefore a key factor in determining the Pf developmental kinetics for multiple parasite strains. Transcriptome analysis of Wnt inhibitor treated hepatocytes indicates that improvement of hepatic metabolism, prevention of EMT and cell division may lead to the positive effects on parasite size and maturation. Together, these findings provide steps forward in understanding the intricate interaction between host and parasite within infected hepatocytes which may benefit rational approaches for future therapeutic interventions.

## Materials and Methods

### Ethics

Primary human liver cells were freshly isolated from remnants of surgical material. The samples were anonymised and general approval for their use was granted in accordance with the Dutch ethical legislation as described in the Medical Research (Human Subjects) Act. It was confirmed by the Committee on Research involving Human Subjects in the region of Arnhem-Nijmegen, the Netherlands. Approval for use of remnant, anonymized surgical material for transcriptome analysis was specifically confirmed by the Committee on Research involving Human Subjects, in the region of Arnhem-Nijmegen, the Netherlands (CMO-2019-5908).

### PHH isolation from liver segments

Primary human hepatocytes were isolated from patients undergoing elective partial hepatectomy as previously described [23, 27]. Freshly isolated hepatocytes are suspended in complete William’s B medium (WB): William’s E with glutamax (ThermoFisher Scientific: 32551087), 1x Insulin/transferrin/selenium (ThermoFisher Scientific: 41400045), 1mM sodium pyruvate (ThermoFisher Scientific: 11360070), 1x MEM Non-essential amino acid solution (ThermoFisher Scientific: 11140035), 100 units/ml Penicillin-Streptomycin (ThermoFisher Scientific: 15140122) and 1.6μM of dexamethasone (Sigma Aldrich: D4902). PHHs were plated at 62,500 cells per well in 96 well format and kept in a 37°C (5% CO2) incubator with daily medium changes 96-well plates (Falcon: 353219) precoated with Type 1 collagen solution from rat tail (Sigma Aldrich: C3867).

### Generation of sporozoites for liver infection

Pf asexual and sexual stages were cultured in a semi-automatic system as described [45–47]. *Anopheles stephensi* mosquitoes were reared at the Radboud University Medical Center Insectary (Nijmegen, the Netherlands) in accordance with standard operating procedures. Salivary glands from infected mosquitoes were hand-dissected and collected in WB medium. Collected glands were homogenized using home-made glass grinders and sporozoites were counted in a Burker-Turk chamber using phase-contrast microscopes. Immediately prior to infection of human hepatocytes, the sporozoites were supplemented with heat-inactivated human sera (HIHS) at 10% of the total volume (i.e. WBH).

### Medium compositions and duration

#### Different plating time after isolation (*Figure 1B*, C)

Isolated PHH were plated in WB medium. The following day, it was changed to WBH: this medium was refreshed daily until the end of the experiment on day 26 post plating. In figure 1D and E, isolated PHH were again plated in WB medium. The following day, it was changed to William’s E with glutamax (ThermoFisher Scientific: 32551087) supplemented with 1x B27^TM^ Supplement (ThermoFisher Scientific: 17504044) and 100 units/ml Penicillin-Streptomycin (ThermoFisher Scientific: 15140122) which was referred to as B27 medium or mock in the article. On the day of infection, B27 medium was removed and replaced with sporozoites suspended in WBH for three hours. After the infection process has occurred (i.e. after three hours), the WBH (sporozoite) medium was replaced with B27 medium until the conclusion of the experiment.

#### Different medium treatments

Isolated PHH were plated in WB medium. The following day, it was changed to the following medium conditions: B27 or B27 supplemented with 20μM of Forskolin (Enzo Life Sciences: BML-CN100-0010), or 10μM of SB431542 (Tocris: 1614), or 0.5μM of IWP2 (Tocris: 3533), or 5μM of DAPT (Tocris: 2634) or 0.1μM of LDN193189 (Tocris: 6053) or the combination of all the compound inhibitors (i.e. 5C). The hepatocytes were kept on this treatment for another 6 days (i.e. 7 days post plating) and then infected with sporozoites where the medium composition changes to WBH for three hours. After three hours, the monolayers were returned to their respective treatments.

#### Investigating the Wnt pathway

Isolated PHH were plated in WB medium. The following day, it was changed to the following medium conditions: B27 or B27 supplemented with 0.5μM of IWP2 or 3μM CHIR-99021 (Sigma-Aldrich: SML1046). The hepatocytes were kept on this treatment for another 6 days (i.e. 7 days post plating) and then infected with sporozoites where the medium composition changes to WBH for three hours. After three hours, the monolayers were returned to their respective treatments.

### Immunofluorescence readout

Monolayers were fixed with 4% paraformaldehyde (ThermoFisher Scientific: 28906) for 10 minutes and permeabilised using 1% Triton for 5 minutes. The samples are stained with the various primary Pf or human antibodies: Rabbit PfHSP70 at 1:75 dilution (StressMarq Biosciences: SPC186), Mouse PfMSP1 at 1:100 dilution (Sanaria and NIH/NIAD: AD233), and mouse human glutamine synthethase at 1:100-250 dilution (Abcam: ab64613). Secondary antibodies were used at these following dilutions: Goat anti-rabbit Alexa Fluor 594 at 1:200 dilution (ThermoFisher Scientific: A11012) and goat anti-mouse Alexa Fluor 488 at 1:200 dilution (ThermoFisher Scientific: A11029). DAPI was used at 300 nM to stain the nuclear material of the monolayer.

### Microscopy

The Zeiss LSM880 with Airyscan at 63x objectives (oil) and 2x zoom were used for detailed images. For high content images, the Zeiss Axio Observer Inverted Microscope Platform with AI assisted experimental startup was used. The images were acquired at 20x objectives with a numerical aperture of 0.8.

### Data analysis using FIJI

#### Infection rate

For each well, 77 images were acquired in a tiled format. Approximately half (i.e. 39) images were counted on FIJI [48] for NF135 and NF175 infections. All the tiles were counted for NF54 due to the lower infection rate. The number of hepatic nuclei were counted for 1% of the total image (i.e. 7-10 images) and then extrapolated to get number of PHHs per well.

#### Measurement of schizont size

See Yang et al for further details [23]. Images obtained on the high content microscope were opened in FIJI. Random images were chosen until at least 100 schizonts were measured (per well) unless the infection is with NF54 (at least 50 schizonts). Schizonts were selected using the region of interest (ROI) tool based on PfHSP70 positivity (red channel) and measured. This ROI mask is applied onto the other colour channels i.e. blue and green to obtain values of nuclear content (DAPI) and hGS or PfMSP1 signal. For hGS and PfMSP1, background non-specific staining was considered and subtracted from the final signal. Further details regarding the methods can be found in [21, 23].

### RNA Isolation and RNA sequencing library preparation

Freshly isolated or in vitro cultured hepatocytes (mock or treatment) were homogenized in TRIzol solution and cryopreserved at −80°C. RNA was subsequently isolated using the Zymo research Direct zol RNA purification kit (Cat. No. #R2053). The isolated RNA was assessed for quality and quantified using Nanodrop and agarose gel electrophoresis. RNA sequencing libraries from isolated/purified RNA were prepared using the Kapa mRNA Hyperprep Kit (Cat. No. #KK8581) using slight modifications of the manufacturer’s instructions for mRNA enrichment method. The input total RNA for the various libraries was selected between 150ng, 250ng or 500ng. The enriched mRNA was fragmented at 94°C for 6min and dA tailing performed at 55°C for 15min. In order to better capture the AT rich *Pf* transcriptome we modified the PCR protocol as follows: Initial denaturation at 98°C for 2min; Cycling denaturation at 98°C for 20sec, annealing + extension at 62°C for 2min; final extension at 62°C for 3min. We selected PCR cycles for library amplification (12, 11 and 10 respectively) based on starting input total RNA amount. The amplified libraries were quantified using Denovix dsDNA high sensitivity kit (Cat. No. #TN145)on a Qubit fluorometer. Further qualitative assessment of library size distribution was performed using the Agilent high sensitivity DNA kit (Cat. No. #5067-4626) on the Agilent 2100 bioanalyzer platform. Each library was sequenced in 42bp paired-end format for roughly 18 million reads on the Illumina Nextseq 500 platform.

### RNA sequencing data analysis

The sequencing endline fastq files were quality checked using FASTQC (ver. 0.11.9). The reads were subsequently trimmed for quality (-q 30) and adapter removal using the Trim_Galore software (ver. 0.6.7). The trimmed reads were aligned onto the *Homo sapiens* ver. 38 genome using STAR aligner (ver. 2.7.10a). The reads mapped onto the respective genomes were counted using the –quantMode GeneCounts option in STAR. Reads mapping only to sense strand were corrected for batch effects stemming from different hepatocyte donors using the Combat-seq tool (sva package ver. 3.44.0). The batch corrected counts were used for subsequent differential expression analysis (for conditions/treatments) in DESEq2 package (ver. 1.36.0) in RStudio (ver. 4.2.2 “Innocent and Trusting”). PCA plot for the various sequenced libraries/samples was generated using plotPCA command. Volcano plots for differentially expressed genes and genes of interest were generated using custom scripts and ggplot2 (ver. 3.4.2) in R. Gene Ontology enrichment analysis was performed in using the clusterProfiler tool (ver. 4.4.4) with subsequent dot-plots generated using inbuilt functions in ggplot2. Heatmaps for differentially expressed genes were generated using the Morpheus online tool by Broad Institute.

### Statistical analysis

For the majority of the experiments, at least three biological replicates were performed with two technical replicates. All statistical tests were performed using Prism 10. See figure legends for details regarding the statistical tests.

## Data availability

The next-generation sequencing dataset associated with this study has been uploaded to Gene Expression Omnibus: GSE263643 The dataset comprises of: The metadata sheet with information on experiments and samples Raw data files: fastq files for RNA-sequencing Processed data files: i) raw counts files generated from Combat-seq and ii) normalized count files generated from DESeq2

## Author contributions

AK, GJG, HW, AS, and ASPY performed the experiments. JHWdW coordinated the collection of the fresh human liver segments. ASPY and AK performed the analysis on the data. TB was involved in the writing and reading of the manuscript. AK, HW, RWS, RB and ASPY were involved in the conceptualization and writing of the manuscript.

## Acknowledgements

We are grateful for R. Stoter, W. Graumans, M. Vegte-Bolmer, A. Pouwelsen, L. Pelser-Posthumus and J. Kuhnen of the Malaria and Insectary Unit at the Radboud University Medical Center for parasite, mosquito and sporozoite production. We would like to thank the Microscopic Imaging Center (MIC) of the Radboud University for access to its facilities. A.S.P.Y was a recipient of the Veni grant from the Dutch Research Council (NWO) talent program (VI.Veni.192.171) and ZonMW Off-Road grant (04510012010050). A.K was supported by the Dutch Research Council (ZonMW-TOP-grant #91218010).

