## Supplementary figures for "Modulation of host signalling pathways reveal a major role for Wnt signalling in the maturation of *Plasmodium falciparum* liver schizonts"

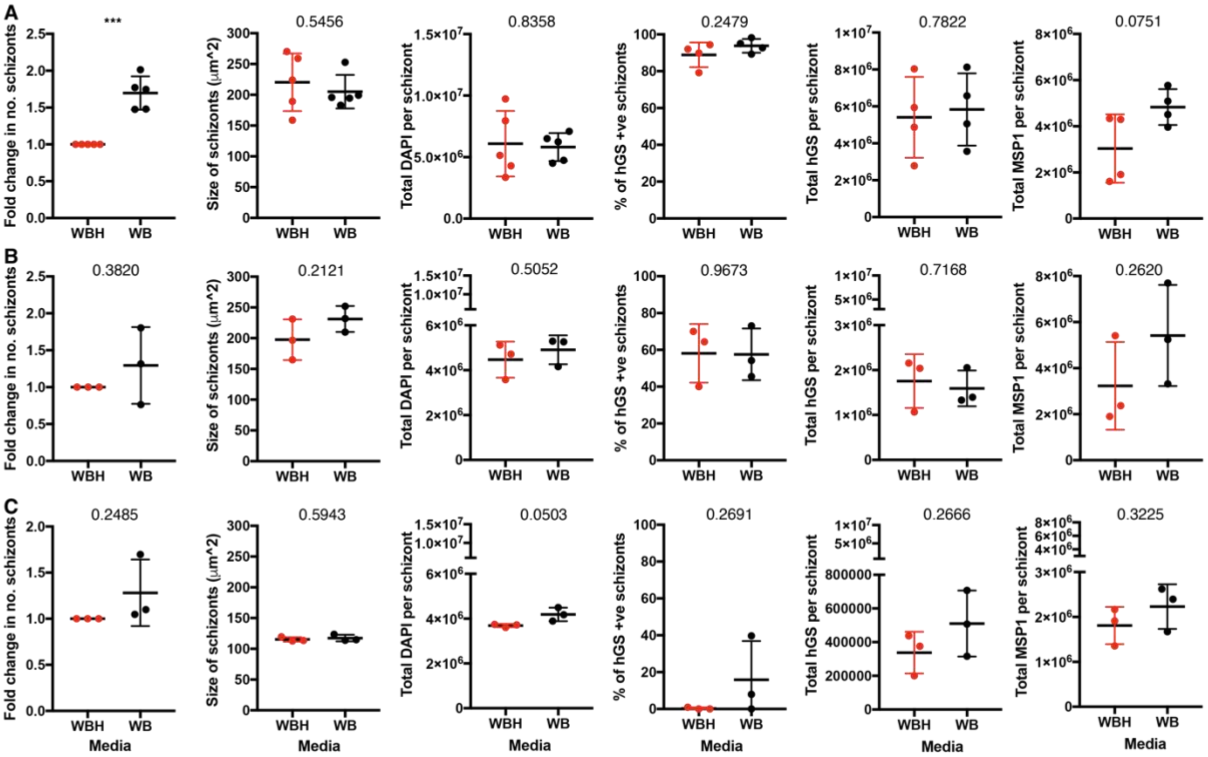


**Figure S1: The impact of heat inactivated human sera on Pf schizont development.** PHH monolayers were cultured in either William's B (WB) or William's B supplemented with 10% heat inactivated sera (WBH) for six days p.p. and infected with either NF175 (B) or NF135 (C) or NF54 (D). Data are shown for respectively schizont number, size, total DAPI content, percentage of human glutamine synthetase (hGS) positive schizonts, total hGS level per schizont and total merozoite surface protein 1 (MSP1) level per schizont. As for the number of schizonts, each dot represents an independent experiment showing the mean of two replicates. As for the size and total DAPI content graphs, each dot represents an independent experiment showing the mean of the median of at least 100 schizonts per replicate (2 replicates) were measured (except in NF54 condition where there is not enough schizonts present). For the percentage of hGS positive schizonts, each dot represents an independent experiment where at least 100 schizonts were examined from one replicate. For the total hGS level and total MSP1 level, each dot represents an independent experiment of the median from at least 100 schizonts (except in NF54 condition where there is not enough schizonts present). The error bars show the mean with the standard deviation. The p-values are generated by performing an unpaired t-test between WBH and WB.

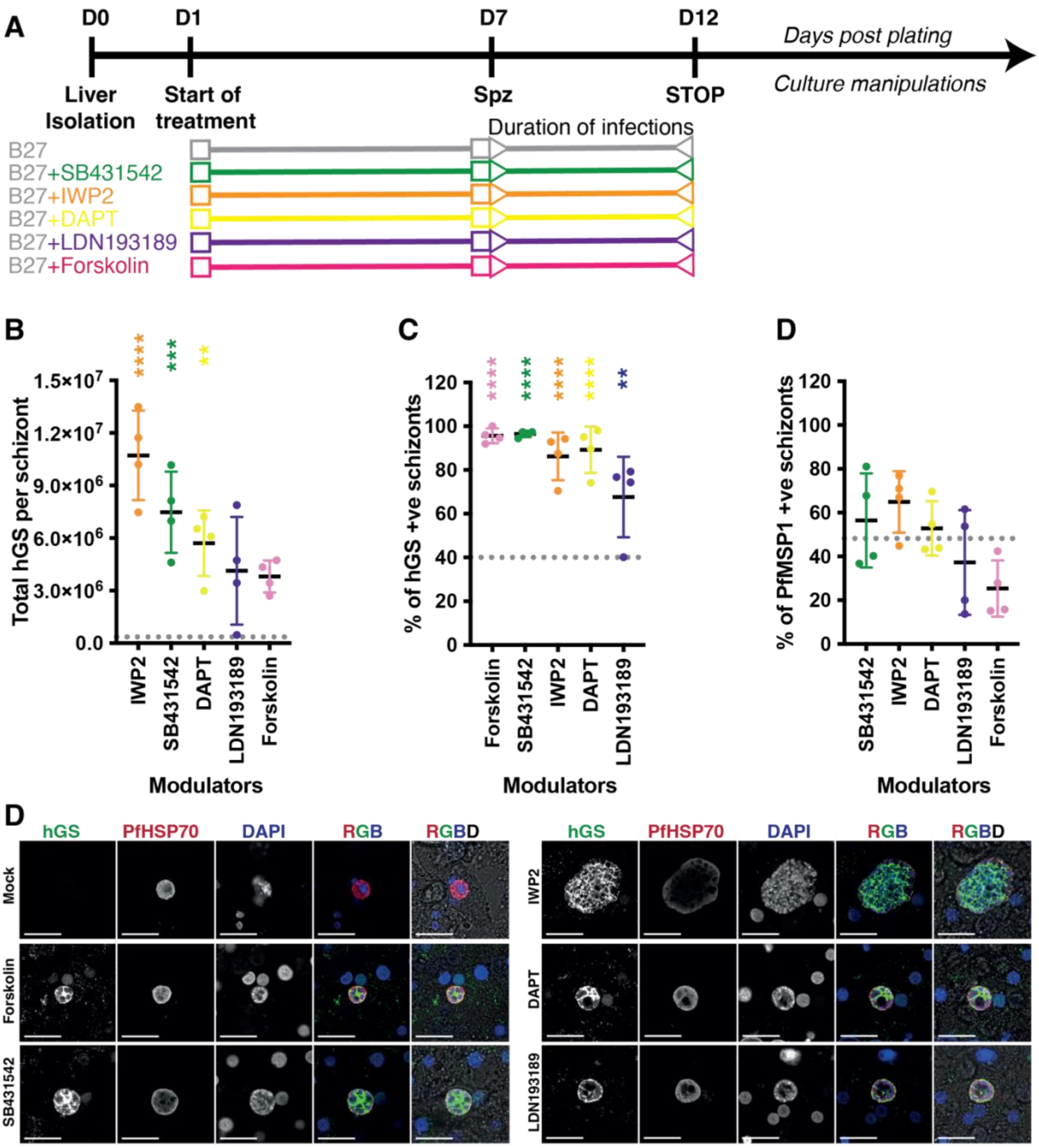


**Figure S2: Effect of specific inhibitors of PHH gene-signalling pathways on the expression Pf schizont markers** A) Schematic experimental setup of Figure 3 and S3. PHHs were treated for 6 days p.p with either B27 or B27 supplemented IWP2/SB431542/Forskolin/DAPT/LDN193189. The experiment was analysed at day 5 p.i.. with NF175 B) Total hGS per schizont with each dot representing the median of at least 100 schizonts. The percentage of hGS (C) and MSP1 (D) positive schizonts: each dot is an independent experiment where at least 100 schizont is measured. The grey dotted line (per graphs B-E) shows the median measurements of schizonts grown in B27 (from four independent experiments) as each host pathway inhibitor treatment was made in media containing B27. The mean and standard deviation is shown for each graph. The p-values from a Dunnett's multiple comparisons test are displayed. D) Representative confocal images (from four independent biological experiments) showing schizonts grown in B27 alone and B27 supplemented with IWP2, SB431542, DAPT, LDN193189, Forskolin stained with hGS, HSP70 and DAPI. Scale bar is 25 microns.

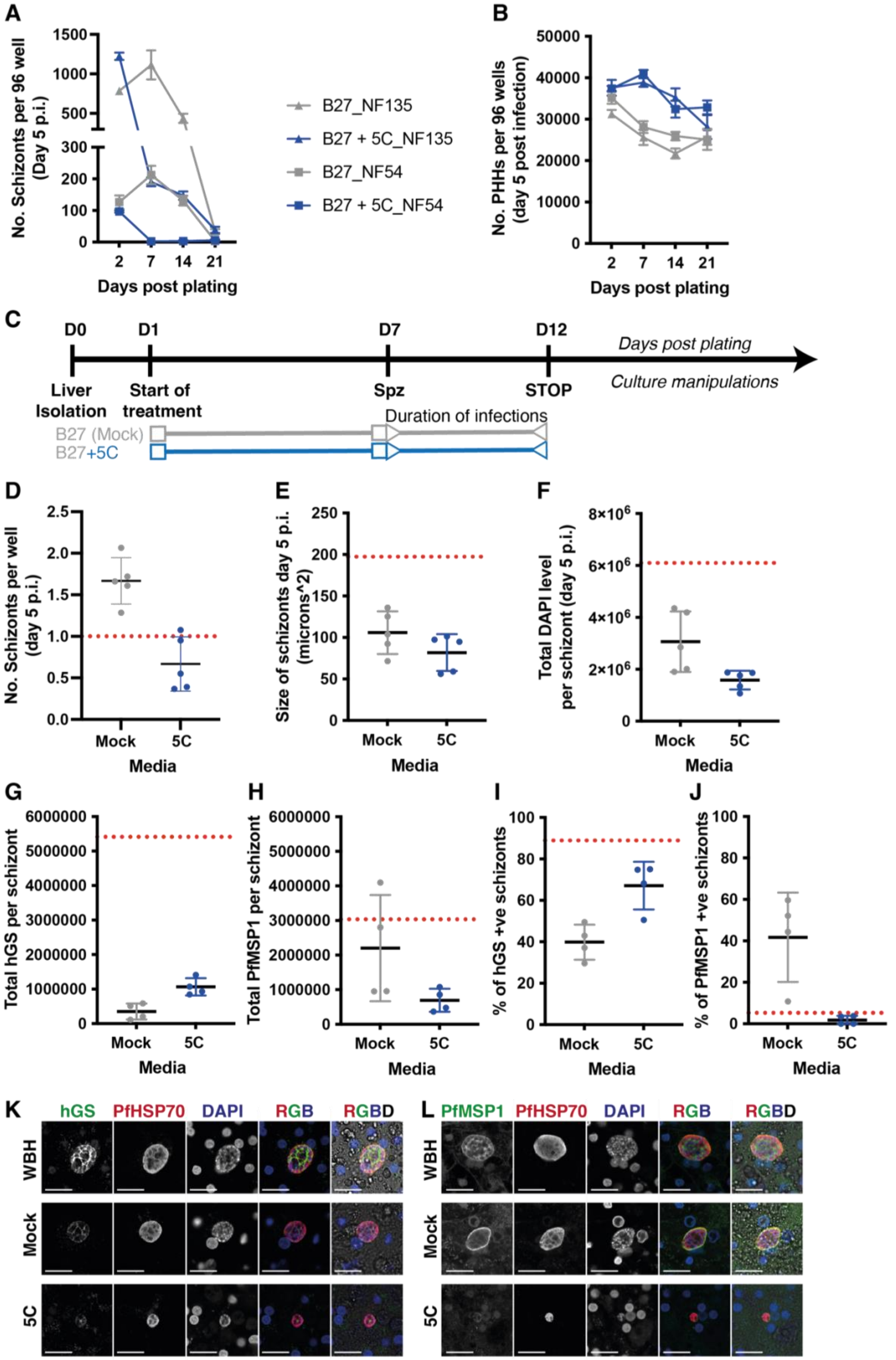
A
No. Schizonts per 96 well (Day 5 p.i.)
Days post plating
B27_NF135
B27 + 5C_NF135
B27_NF54
B27 + 5C_NF54
B
No. PHHs per 96 wells (day 5 post infection)
Days post plating
C
D0
D1
D7
D12
Days post plating
Liver Isolation
Start of treatment
Spz
STOP
Culture manipulations
Duration of infections
B27 (Mock)
B27+5C
D
No. Schizonts per well (day 5 p.i.)
Mock
5C
Media
E
Size of schizonts day 5 p.i. (microns^2)
F
Total DAPI level per schizont (day 5 p.i.)
G
Total hGS per schizont
H
Total PfMSP1 per schizont
I
% of hGS +ve schizonts
J
% of PfMSP1 +ve schizonts
K
hGS
PfHSP70
DAPI
RGB
RGBD
WBH
Mock
5C
L
PfMSP1
PfHSP70
DAPI
RGB
RGBD

**Fig S3: The effect of B27 or B27 supplemented with 5C on *Pf*-permissiveness and schizont development in PHH.** The number of schizonts (A) and hepatocytes (B) on day 5 p.i. for NF135 and NF54 (n=1) at indicated time points p.p. when cultured in either B27 medium (grey) or B27 supplemented with 5C (blue). Each dot shows the mean of three replicates and the error bars show the standard deviation. C) Experimental setup of graphs D-L. PHHs were cultured for 6 days p.p. with either B27 or B27 with 5C and infected with NF175 parasites and analysed at day 5 p.i.. D) The number of schizonts per well was normalised to the number of schizonts counted in WBH (red dotted line). Each dot is the mean of an independent experiment with two replicates. For the schizont size (E) and total DAPI content per schizont (F), each dot represents the mean of an independent experiment of at least 100 schizonts. For the total hGS (G) and MSP1 (H) per schizont, each dot represents the median of an independent experiment of at least 100 schizont For the percentage of hGS (I) and MSP1 (J) positive cells, each dot represents an independent experiment of at least 100 schizonts. D-J shows the mean with the standard deviation). Representative confocal images (from four independent experiments) showing schizonts grown in WBH or B27 alone or B27 supplemented with 5C on day 5 p.i. stained with hGS (K) or MSP1 (L), HSP70 and DAPI. Scale bar is 25 microns.

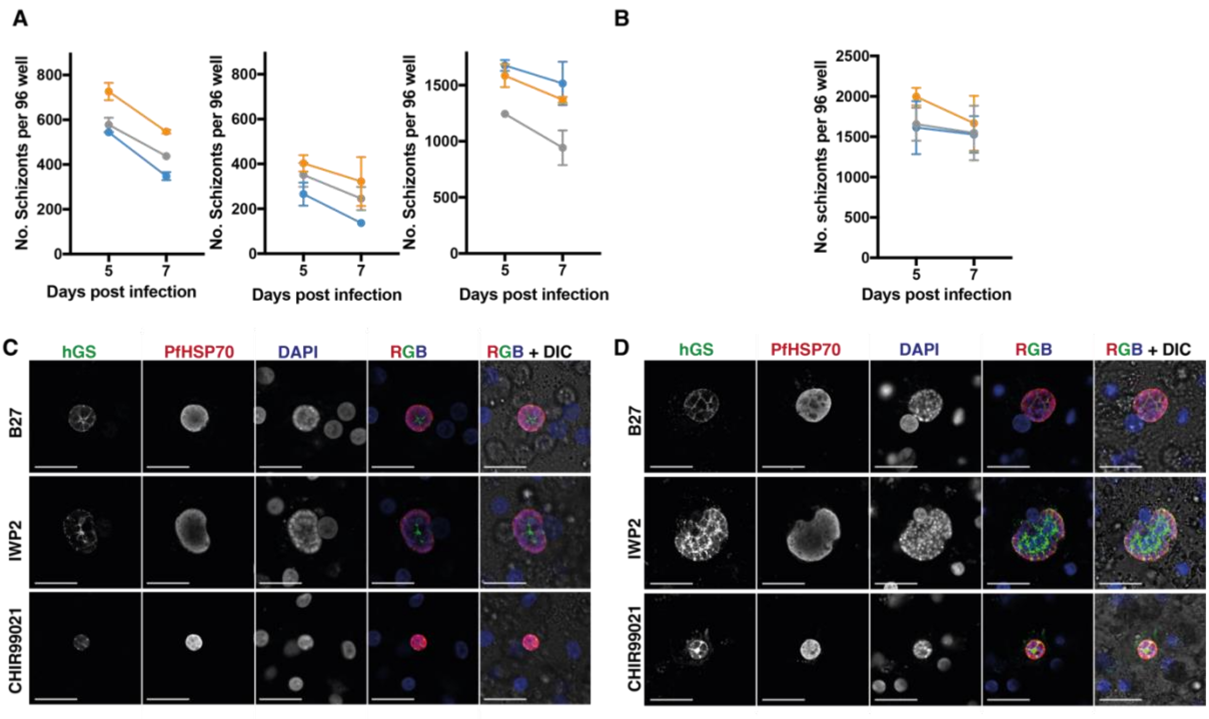


**Figure S4: The impact of the Wnt signalling pathway on numbers of developing schizonts.** A) The number of schizonts in NF135 infected PHHs treated with B27 or B27 supplemented with IWP2/CHIR99021. Each graph shows an independent experiment, each with duplicates per condition. B) Number of schizonts in NF175-infected PHHs treated with B27 or B27 supplemented with IWP2/CHIR99021. Data show mean and the standard deviation of three independent experiments, each with duplicates. Representative confocal images (from three independent biological experiments) showing NF135 (C) and NF175 (D) schizonts B27 or B27 supplemented with IWP2/CHIR99021 on day 5 p.i. stained with hGS, HSP70 and DAPI. Scale bar is 25 microns.
